# Disrupting Pregnane X Receptor Signaling Overcomes Temozolomide Resistance in Glioblastoma via Succisa pratensis–Derived Metabolites

**DOI:** 10.64898/2026.06.22.733681

**Authors:** Francesca Servidio, Fabio Pirovano, Sofia Remedia, Chiara Pellizzer, Martina Nespoli, Bruno Giovanni Galuzzi, Marcella Bonanomi, Sara Mallia, Mauro Commisso, Flavia Guzzo, Clarissa Gervasoni, Daniela Gaglio, Manuela Moriggi, Daniele Capitanio, Gloria Rita Bertoli, Alessandro Giammona, Alessia Lo Dico

**Author notes:** **Corresponding author: Dr Alessia Lo Dico,** via Fratelli Cervi, 93, 20054 Segrate, Milan, Italy. These authors contributed equally to this work (Alessandro Giammona, Alessia Lo Dico).

## Abstract

Glioblastoma remains a highly aggressive and therapy-resistant brain tumor, with limited benefit from the current standard-of-care regimen combining surgery, radiotherapy, and temozolomide. Overcoming chemoresistance therefore represents a critical unmet clinical need.

Here, we investigate the anticancer potential of *Succisa pratensis* and its ability to enhance TMZ efficacy in GBM models. Treatment with *S. pratensis* markedly reduced cell proliferation and migration while significantly increasing sensitivity to TMZ. Integrated multi-omics analyses revealed extensive metabolic rewiring, characterized by suppression of central carbon metabolism and activation of stress-adaptive pathways.

Mechanistically, we identify the Pregnane X Receptor, a key regulator of drug metabolism and chemoresistance, as a central node affected by treatment. Although *S. pratensis* increased PXR expression, this was not accompanied by induction of canonical downstream targets, including *MDR1* and *ALDH1A1*, indicating a functional impairment of PXR transcriptional activity. Consistently, pharmacological inhibition of PXR using the antagonist SPA70 further potentiated the cytotoxic effects of *S. pratensis* and TMZ.

Docking analyses suggest that specific secondary metabolites, including apigenin-derived compounds, may interact with the PXR ligand-binding domain, providing a potential molecular basis for this effect.

Collectively, our findings indicate that *S. pratensis* enhances TMZ efficacy by inducing metabolic vulnerability and functionally impairing PXR signaling. These results highlight the therapeutic potential of plant-derived metabolites as adjuvant strategies to overcome chemoresistance in glioblastoma.

**Article Highlights:**

- Succisa pratensis enhances temozolomide efficacy in glioblastoma by reducing proliferation, migration, and clonogenic growth.
- Integrated proteomic and metabolomic analyses reveal extensive metabolic rewiring, with suppression of central carbon metabolism and induction of stress-adaptive pathways.
- Pregnane X Receptor (PXR), a key regulator of chemoresistance, is functionally impaired despite increased expression, resulting in reduced activation of drug-resistance genes.
- Pharmacological inhibition of PXR further potentiates the antitumor effects of Succisa pratensis and temozolomide, promoting apoptotic cell death.
- Apigenin-derived metabolites show high affinity for the PXR ligand-binding domain and emerge as promising candidates to overcome temozolomide resistance in glioblastoma.

## Introduction

Glioblastoma multiforme (GBM) accounts for nearly 80% of malignant primary brain tumors and remains highly lethal despite therapeutic advances. Median survival is only 14.6 months, with recurrence typically occurring within 7 months. [1–3] Standard treatments, including surgery, radiotherapy, and chemotherapy, offer limited efficacy, and few trials have defined optimal protocols. Temozolomide (TMZ), the gold standard in the STUPP protocol, remains central to therapy, while additional strategies such as nitrosoureas, bevacizumab, tyrosine kinase inhibitors, and immunotherapy are being explored to improve outcomes. [4–6]

Drug resistance poses a major challenge in GBM treatment, It involves various biological and clinical factors, such as intrinsic resistance due to the tumor’s heterogeneity, the physical blood-brain barrier (BBB), which limits drug penetration into the brain, or the overexpression of efflux transporters, such as of ATP-binding cassette (ABC) transporters, that reduce drug efficacy by expelling chemotherapeutics from cells. [7–9]

TMZ efficacy in GBM is hindered by ABC efflux transporters, especially P-glycoprotein (*ABCB1*) and *ABCD1*, which expel the drug from cancer cells and across the BBB, causing chemoresistance. Recent studies have shown that pharmacological inhibition or genetic silencing of these transporters can enhance TMZ sensitivity both in vitro and in vivo [10,11] Biodiversity represents a valuable source of bioactive molecules, and more than 60% of current anticancer drugs derive from natural products. [12] [13]

Previous work by Giammona et al. demonstrated that *S. pratensis* significantly reduces GBM proliferation and migration, promotes apoptosis and mesenchymal-to-epithelial transition (MET), induces extensive metabolic rewiring, and impairs antioxidant defenses through modulation of inflammatory signaling. LC-MS analysis identified several candidate bioactive metabolites, including the flavonoids luteolin-C-hexosyl-O-hexoside and apigenin-C-hexosyl-O-hexoside, the phenolics chlorogenic acid and esculetin, and the saponin Akebia saponin D. [14]

This paper highlights the anti-cancer properties of *S. Pratensis,* which with leaves and flowers containing phenolic acids, flavonoids, and essential oils. Extracts and compounds like luteolin 7-glucoside and apigenin 7-glucoside show antioxidant, antimicrobial, and anticancer effects, modulating inflammatory pathways such as NF-κB in HepG2 liver cancer cells. [15]

The Pregnane X Receptor (PXR) protein has been implicated in GBM chemoresistance, as its activation regulates the expression of drug-metabolizing enzymes and efflux transporters that can reduce TMZ efficacy [16]. PXR is a nuclear receptor involved in xenobiotic sensing and regulation of genes associated with chemoresistance, including CYP3A4, ALDH1A1, ABCG2, and MDR1. High PXR expression correlates with poor prognosis in GBM and several other cancers. [17] [18]

The study’s primary objective is to characterize secondary metabolites’ effects in glioblastoma by elucidating the molecular mechanisms underlying their activity.

Therefore, this study aimed to investigate the role of S. pratensis-derived metabolites in TMZ sensitization and to determine whether modulation of PXR signaling contributes to overcoming GBM chemoresistance.

## Materials and methods

### GBM Cell lines and treatment

U118, LN18, and T98 cells purchased from ATCC and maintained in Dulbecco’s modified Eagle’s medium (DMEM) (Euroclone, Milan, Italy) or RPMI (Euroclone, Milan, Italy) containing 10% heat-inactivated fetal bovine serum (FBS), 50 IU/ml of penicillin and streptomycin, 2 mM glutamine (Euroclone, Milan, Italy) in a humidified atmosphere at 5% CO2 incubator at 37 °C. GBM cells were treated for 24 hours (h) with *S. Pratensis* at 5 mg/ml, Akebia saponin D (Selleck Chemicals Catalog No. S5455), or Apigenin-C-hexosyl-O-(caffeoyl) hexoside (MedChemExpress Cat. No.: HY-N5083) at 100 μM. The double treatment is obtained by adding Temozolomide (TMZ) at 100 μM (Sigma-Aldrich, Germany-Cat. No. T2577).

### Proliferation and wound healing assays

For the proliferation assay, 5 × 10⁴ cells/well were seeded in 24-well plates and treated with biomolecules, secondary metabolites, and/or TMZ. After 24 h, cells were counted using a semi-automated cell counter (Diatech Lab Line, Jesi, Italy). Data are presented as cell viability (fold change relative to control) and cell death (%).

For the wound healing assay, GBM cells (5–8 × 10⁴ cells/well) were seeded in 12-well plates and cultured to 70–80% confluence. A linear scratch was generated using a 200 μl pipette tip, detached cells were removed by washing, and fresh medium containing the biomolecules was added. Images were acquired after 24 h using a phase-contrast microscope (4× and 10×). Wound area was quantified with ImageJ software, and cell migration was expressed as percentage wound closure according to the formula: wound closure = [(A₀ − A₂₄)/A₀] × 100, where A₀ is the wound area immediately after scratching and A₂₄ is the wound area after 24 h.

### Cell cycle

A total of 1 × 10⁶ harvested cells were fixed in cold ethanol by dropwise addition under vortex agitation. For cell cycle analysis, cells were washed twice with PBS containing 2% FBS and resuspended in 1 mL of staining buffer (PBS, 2 mM EDTA, 0.6% trypsin, 2% FBS, 0.2% NP-40, and 15 µg/mL propidium iodide). After 30 min incubation at room temperature in the dark, samples were acquired using a FACS Celesta flow cytometer (BD Life Sciences, New Jersey, USA). Data were analyzed with FlowJo v10 using the Watson pragmatic model to determine the distribution of cells in G0/G1, S, and G2/M phases.

### Immunofluorescence

For IF analysis, 2 × 10⁴ GBM cells were seeded on glass coverslips and treated with biomolecules after 24 h. Following an additional 24 h, cells were washed with PBS, fixed with 4% paraformaldehyde for 30 min at 37 °C, permeabilized with PBS/5% FBS/0.3% Triton X-100 for 10 min at 4 °C, and blocked with 2% BSA in PBS/0.1% Tween-20 for 30 min at room temperature. Cells were incubated overnight at 4 °C with anti-human PXR antibody (sc-48340, Santa Cruz), followed by Alexa Fluor™ 488-conjugated secondary antibodies (Invitrogen) for 30 min at room temperature in the dark. Nuclei were counterstained with DAPI (Thermo Fisher Scientific). Images were acquired with a Leica STELLARIS 5 microscope (63× oil objective) and analyses performed using ImageJ software (NIH, USA).

### Quantitative real-time PCR

Total RNA was extracted using TRIzol reagent (Zymo Research) and reverse-transcribed into cDNA with PrimeScript RT Master Mix (Takara). Quantitative real-time PCR was performed in triplicate using SYBR Green chemistry (2X Green Fast qPCR Mix, Fisher Molecular Biology, FMB). Primer sequences (Eurofins, Milan, Italy) are listed in Table S1. Relative gene expression was normalized to β-ACTIN and calculated using the ΔΔCt method (Livak & Schmittgen, 2001), with results expressed as fold induction relative to vehicle-treated controls.

### Survival Analyses

Overall survival data were obtained from the GEPIA database (http://gepia.cancer-pku.cn/) and analyzed using the log-rank (Mantel–Cox) test, presented as a Kaplan–Meier survival curve

### Statistical Data Analysis

Experiments were performed in triplicate, and results are presented as mean ± SD. Statistical analyses were conducted using t-tests or one-/two-way ANOVA followed by Dunnett’s or Bonferroni’s post hoc tests (GraphPad Prism 4).

Metabolomic data were analyzed with Mass Profiler Professional 15.1 (Agilent Technologies). Raw data were log₂-transformed, Pareto-scaled, and filtered to retain entities detected in 100% of samples in at least one condition. Statistical significance was assessed by one-way ANOVA (p < 0.05) with Benjamini–Hochberg FDR correction. Significant metabolites were visualized by hierarchical clustering.

## Results

### S. pratensis induces a coordinated metabolic collapse and stress-adaptive rewiring in glioblastoma cells

To define the metabolic impact of S. pratensis treatment in GBM cells, we performed an integrated multi-omics analysis combining quantitative proteomics and untargeted metabolomics.

Proteome analysis, performed using liquid chromatography coupled with electrospray ionization tandem mass spectrometry (LC-ESI-MS/MS) and TMT isobaric labeling, identified 965 proteins differentially expressed compared to the DMSO-treated control, as determined by ANOVA followed by Tukey’s post hoc test with FDR correction (p-value < 0.05). The identification data for the differentially expressed proteins (DEPs) are available at https://doi.org/10.13130/RD_UNIMI/ME5M6X.

Ingenuity Pathway Analysis (IPA) of the enriched “Canonical Pathways” in S. pratensis-treated cells revealed the inhibition of 211 pathways involved in cell cycle regulation, energy metabolism, protein synthesis, nucleic acid synthesis, DNA repair processes, and detoxification mechanisms (e.g., Detoxification of ROS, HIF1α signaling, KEAP1-NFE2L2 pathway, NRF2-mediated oxidative stress response, and xenobiotic metabolism via PXR signaling).

In S. pratensis-treated cells, Rho family GTPase signaling (RHO GTPase cycle, RHOA, ILK) was inhibited, impairing cytoskeleton dynamics, adhesion, motility, proliferation, and differentiation, while the inhibitory RHOGDI pathway was activated. Conversely, Ephrin-A and Granzyme-A signaling were upregulated, suggesting enhanced tumor-suppressive activity and increased clearance of transformed cells (Fig. 1 and S1A) [29].

**Figure 1:**
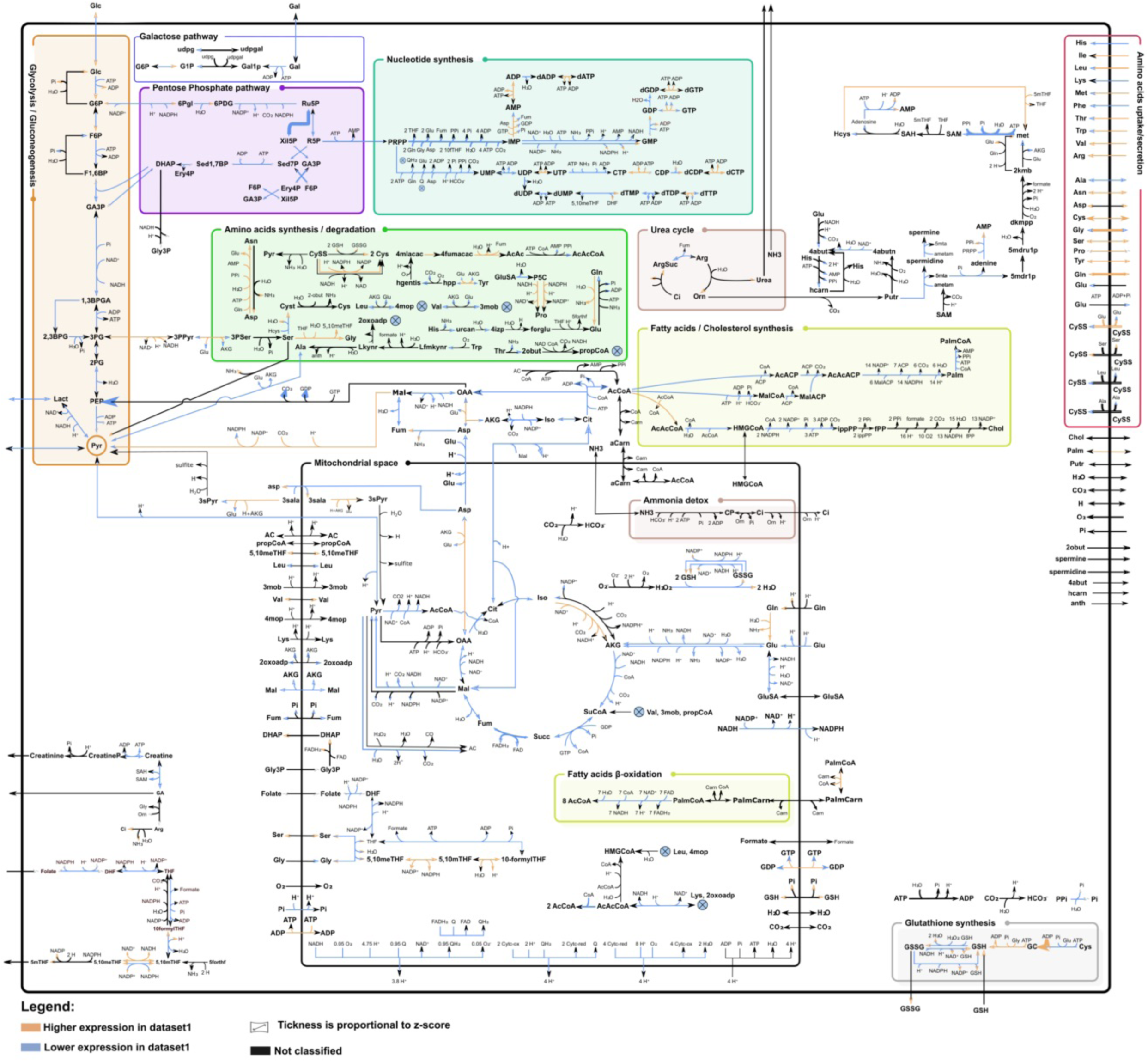
*S. Pratensis* treatment profoundly disrupts glioblastoma cell metabolism. The ENGRO2 metabolic network map visualizes RPS and RAS differences between the *S. Pratensis* treatment and the control condition. Proteomic data are represented by arrows, with orange arrows indicating higher scores and blue arrows indicating lower scores. Metabolomic data are represented by arrowheads, with orange arrowheads indicating higher scores and blue arrowheads indicating lower scores. Arrows or arrowheads shown in black indicate missing information.

Untargeted mass spectrometry–based metabolomics analysis revealed that S. pratensis broadly suppresses central carbon metabolism in U118 GBM cells (Fig. S1B). Key intermediates of glycolysis, the TCA cycle and the pentose phosphate pathway were markedly reduced, together with metabolites involved in ketone body metabolism, fatty acid transport and creatine metabolism

Amino acids associated with one-carbon metabolism, branched-chain amino acid metabolism and glutamate metabolism were broadly decreased.

moIn parallel, nucleotides and their precursors were consistently decreased, including pyrimidine intermediates (UMP, dCMP, UDP, UDP-glucose, UDP-glucuronate, GDP-glucose) and purine intermediates (adenylsuccinic acid, hypoxanthine). Additional reductions were observed in collagen- and tyrosine-related metabolites (hydroxyproline, 4-hydroxyphenylpyruvate), the uronic acid pathway (gluconate, UDP-glucuronate), and the mevalonate/isoprenoid pathway (geranyl diphosphate).

In contrast, several stress-adaptive and salvage pathways were upregulated. Metabolites associated with glutathione metabolism showed consistent increases, including GSH, oxidized GSH, L-cysteine, and γ-glutamylcysteine. Folate one-carbon intermediates (5,10-methenyl-THF and 5,10-methylene-THF) also increased, together with purine salvage metabolites (AMP, GMP, IMP, inosine, adenine, and adenosine) and pyrimidine intermediates (cytidine, dCDP).

Levels of key nutrients (glucose, glutamine), antioxidant molecules (hypotaurine, carnosine), and amino acids linked to nutrient stress responses (aspartate, arginine, asparagine, and tyrosine) were elevated. Glycerol-3-phosphate and mevalonic acid also increased, suggesting a rerouting of carbon flux toward lipid remodeling and a bottleneck in isoprenoid biosynthesis. Altogether, these data indicate that S. pratensis treatment suppresses energy-producing and anabolic pathways while enhancing redox defense, one-carbon metabolism, and nucleotide salvage, consistent with a coordinated metabolic stress response (Fig. S1B).

To obtain a comprehensive overview of these alterations, we integrated proteomics and metabolomics data into the ENGRO2 metabolic network [28] using the Marea4Galaxy tool [27]. From metabolomics, we calculated Reaction Propensity Scores (RPS), obtaining 256 values corresponding to reactions with available data. From proteomics, we derived Reaction Activity Scores (RAS), resulting in 237 values. These were projected onto the metabolic network using the MAREA module to enable integrated visualization (Fig. 1).

This analysis revealed a general downregulation of central metabolic pathways at the protein level, consistent with the IPA results, including glycolysis, the pentose phosphate pathway (PPP), the TCA cycle, oxidative phosphorylation, and fatty acid and cholesterol biosynthesis. The only pathways showing significant upregulation were those involved in amino acid uptake, synthesis, and degradation.

At the metabolite level, a concordant downregulation of glycolysis and the TCA cycle was observed, reinforcing the trends seen in the proteomic data. Overall, this integrated metabolic profile indicates a broad suppression of core metabolic activities in S. pratensis-treated cells, likely reflecting reduced proliferative capacity. Amino acid metabolism appears to be the main upregulated pathway, possibly acting as a compensatory mechanism.

Interestingly, glutathione (GSH) biosynthesis showed a divergent behavior, with metabolomics indicating upregulation while proteomics suggested downregulation, pointing to complex regulatory mechanisms governing redox homeostasis.

Our previous studies showed that S. pratensis induces oxidative stress in GBM cells, as evidenced by increased ROS, decreased GSH/GSSG ratio, and downregulation of antioxidant genes. Consistently, metabolic profiling revealed broad pathway rewiring, including increased levels of glucose, glutamine, and antioxidant-related metabolites, with folate metabolism upregulation correlating with enhanced TMZ sensitivity and improved chemotherapeutic efficacy.

### Combined treatment with *S. Pratensis* and TMZ reduces GBM cell proliferation and migratory capacity

To investigate the therapeutic potential of S. pratensis, we evaluated its phytoextract in combination with TMZ on GBM proliferation, cell cycle progression, and migration. Previous studies showed that S. pratensis extract alone markedly reduced GBM aggressiveness without affecting non-cancerous cells. [14] However, its ability to enhance TMZ sensitivity, improving chemotherapeutic efficacy, has not yet been explored. Quantitative analyses using semiautomated cell counting after 24 hours and colony-forming assay after long-term combined treatment with TMZ and S. pratensis revealed a significant decrease in viable GBM cell numbers, accompanied by a concomitant increase in cell death compared with untreated or TMZ-only controls (**Fig. 2A, S2A and S2B**). Cell cycle analysis showed that combined treatment induced G0/G1 accumulation, indicative of G1 checkpoint activation, across all GBM cell lines compared with TMZ alone (**Fig. 2B).** Beyond the effects on proliferation, the combination treatment also markedly reduced the migratory capacity of residual GBM cells, as demonstrated by wound-healing assays (**Fig. 2C**). Immunofluorescence analysis showed that combined treatment markedly disrupted cytoskeletal organization, leading to a pronounced decrease in tubulin fiber density (**Fig. 2D**).

**Figure 2:**
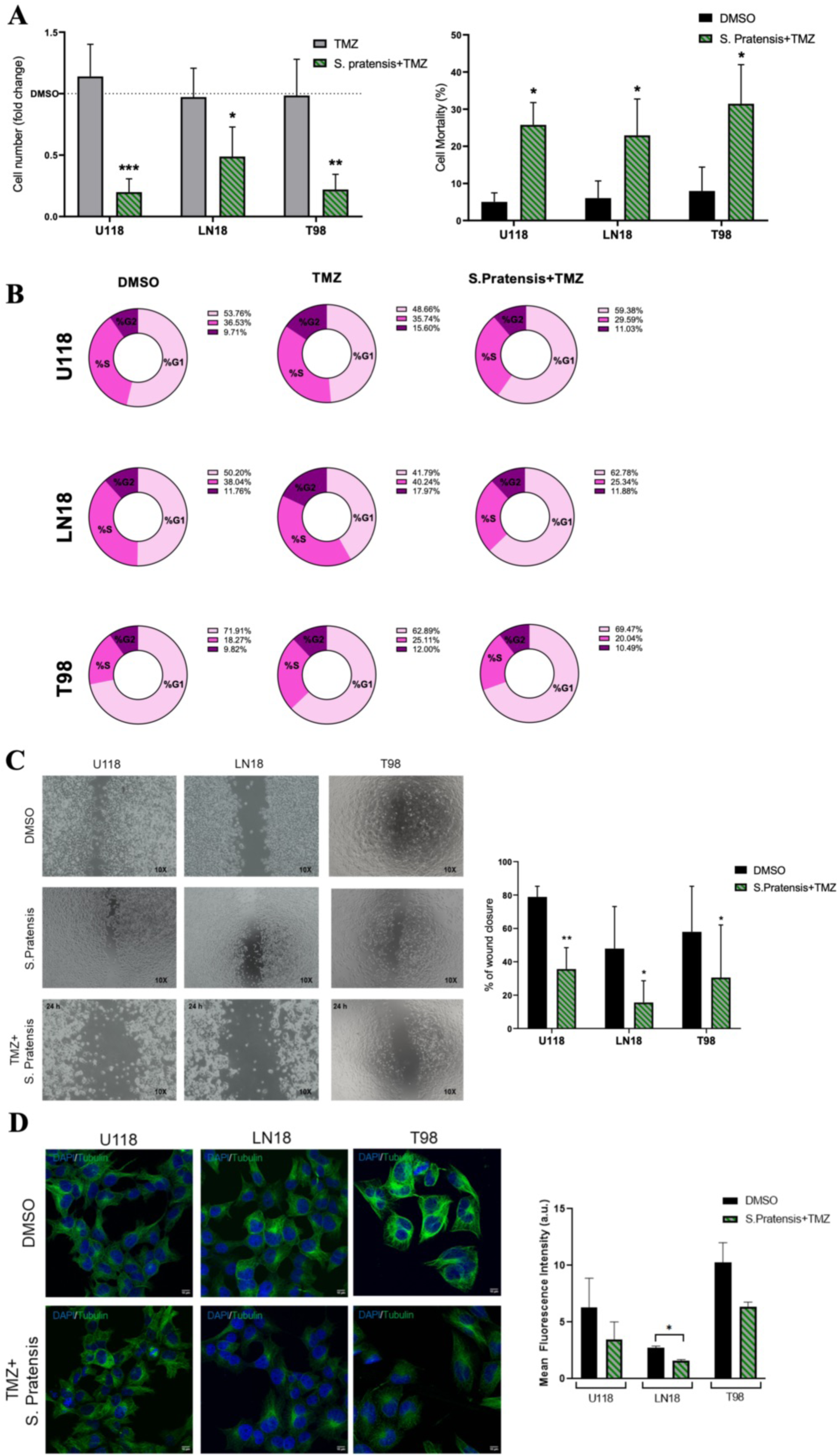
S. Pratensis extract combined with TMZ reduces proliferation and migratory capacity in GBM cells. **(A, left side)** Quantification of cell proliferation, expressed as fold change in cell number relative to the control. Data are presented as mean ± standard deviation (SD) from three independent experiments (N=3). Statistical significance was determined using a t-test (**p < 0.01, ***p < 0.001). **(A, right side)** Cell death was assessed in GBM cells using the Trypan Blue exclusion assay after 24 hours of treatment with DMSO (control) or *S. Pratensis* extract combined with TMZ. Cell mortality is expressed as a percentage of total cells. Data are shown as mean ± standard deviation (SD) from three independent experiments. Statistical significance was determined using a t-test (*p < 0.05). (B) Propidium iodide cell cycle assay performed by flow cytometry analysis on all the GBM cells treated with combined treatment or TMZ alone. **(C, left side)** Representative images of wound closure in GBM cells treated with DMSO (control) or *S. Pratensis* extract combined with TMZ after 24 hours. (**C, right side**) Quantification of wound closure percentage, representing cell migration ability. Data are presented as mean ± standard deviation (SD) from three independent experiments (N=3). Statistical significance was determined using a t-test (**p < 0.01). (**D, left side**) Representative immunofluorescence images of GBM cells cultured for 24 h with double treatment and showing a cytoskeleton rearrangement. The cells were stained with Tubulin antibody, and the Nuclei were counterstained with DAPI. (**D, right side**) Fluorescence relative quantification of Tubulin levels by using ImageJ software.

### Combined Treatment Induces Synergistic Effects Compared to Single Treatments in GBM cells

To evaluate the metabolic disruption induced by the combined treatment with *S. Pratensis* extract and TMZ, we calculated the Reaction Activity Scores (RAS) using the proteomics data. The comparative analysis between the combined *S. Pratensis* and TMZ treatment and the control is shown in **Figure 3**. RAS analysis confirmed a broad suppression of glycolysis, the PPP, TCA cycle, nucleotide synthesis, folate metabolism, and glutathione metabolism in double-treated cells compared with controls. This effect was more pronounced than that observed with S. pratensis alone, suggesting a reduced capacity to sustain DNA synthesis and proliferation. To confirm these observations, we examined the proteomic profile in more detail.

**Figure 3:**
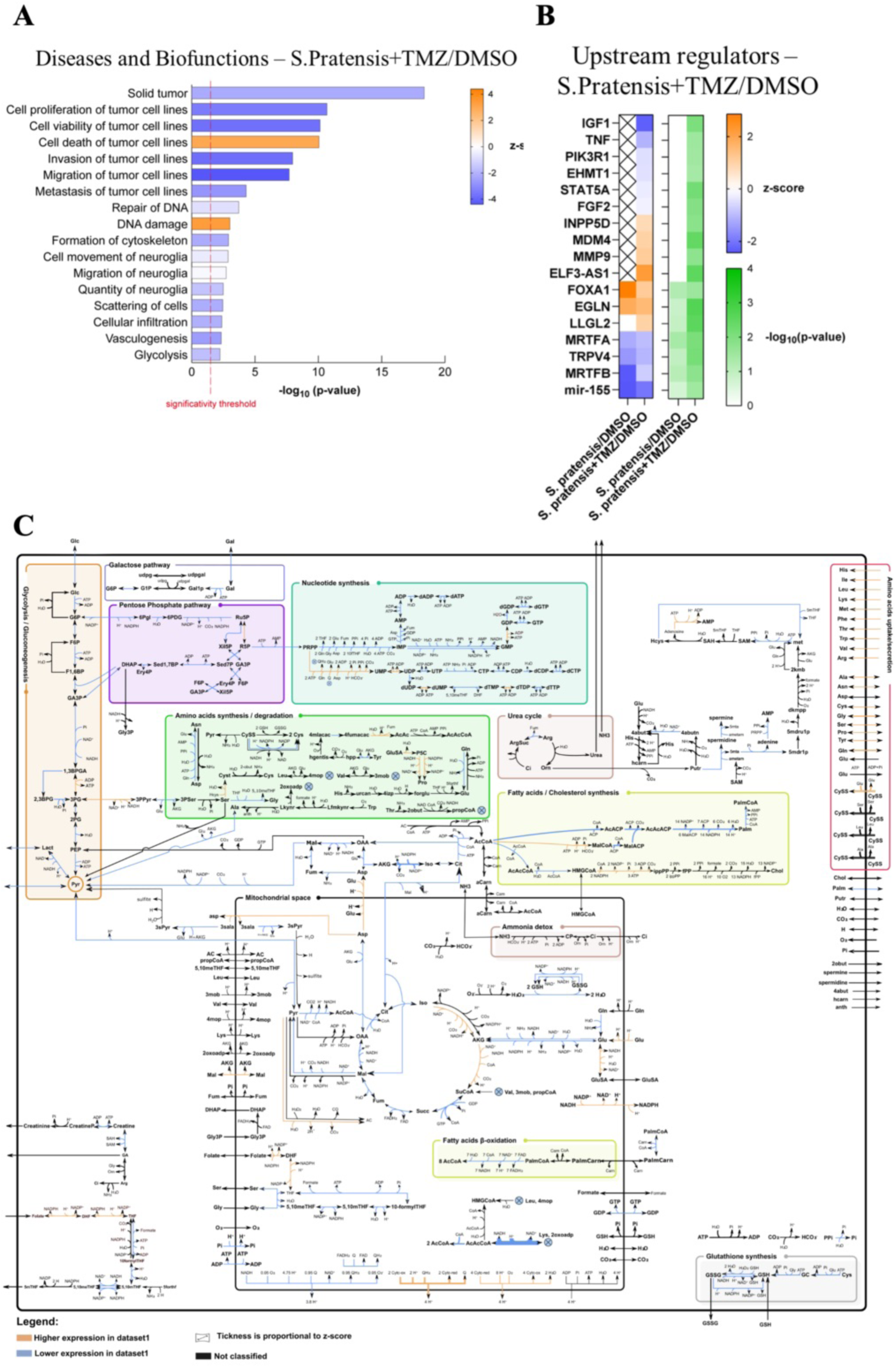
Combined Treatment Induces Synergistic Effects Compared to Single Treatments in GBM cells. **(A)** IPA analysis showing the overrepresented diseases and biological functions in the *S. Pratensis* and TMZ co-treated cell line compared to the control, ranked by p-value. The red vertical line indicates the significance threshold (p = 0.05). Orange bars represent predicted pathway activation, while blue bars represent predicted inhibition, according to the z-score statistic. (**B)** IPA comparison analysis illustrating the cascade of upstream transcriptional regulators that may account for the observed protein expression changes in *S. Pratensis*-treated and co-treated cells compared to the control. The color scale indicates predicted activation (orange) or inhibition (blue) based on the z-score, while green highlights statistically significant regulators. **(C)** The ENGRO2 metabolic network map visualizes RAS differences between the combined treatment and the control condition. Proteomic data are represented by arrows, with orange arrows indicating higher scores and blue arrows indicating lower scores. Arrows or arrowheads shown in black indicate missing information.

Proteomic analysis of the combined treatment with *S. Pratensis* extract and temozolomide (TMZ) revealed 239 differentially expressed proteins compared to the DMSO-treated control, including 65 upregulated and 174 downregulated proteins (ANOVA followed by Tukey’s post hoc test with FDR correction, p-value < 0.05) (https://doi.org/10.13130/RD_UNIMI/ME5M6X). Based on the expression profiles of these proteins, IPA predicted inhibition of proliferation, migration, invasion, metastasis, cytoskeleton organization, glycolysis, and vasculogenesis, together with activation of DNA damage and tumor cell death pathways (**Fig. 3A**).

Upstream regulator analysis identified inhibition of multiple regulators involved in proliferation, migration, inflammation, and survival pathways, consistent with enhanced chemosensitivity and reduced tumor aggressiveness. [30].

*INPP5D,* a key regulator of autophagy maintenance and inflammasome activity, was predicted to be activated together with *EGLN* and *LLGL2*, two tumor suppressor genes associated with cancer progression. [31–34] All these regulators were significantly predicted to be either activated or inhibited in double-treated cells compared with controls. In contrast, they were absent in TMZ-treated cells and either lacking or not significantly predicted in *S. Pratensis*-treated cells.

Nonetheless, *MDM4, MMP9, FOXA1*, and the lncRNA *ELF3-AS1* genes previously reported to be expressed in glioma [35] [11] [36] remained activated, suggesting the persistence of glioma-associated features in double-treated cells. (**Fig. 3B**)

### *S. Pratensis* treatment impairs GBM drug resistance by modulating PXR activity and promoting cell death pathways

To elucidate the mechanisms underlying the reduced growth of TMZ-resistant GBM cells following combined treatment with *S. Pratensis* extract and TMZ, we speculated that a lack of PXR receptor activity and its inability to efficiently promote drug resistance modulation may be involved. This nuclear receptor detects xenobiotics and drugs, thereby should be a promising candidate to justify the diminishing treatment efficacy by upregulating genes related to drug efflux and metabolism (**Fig. 4A**). Therefore, using an in-silico approach, we first examined the overall survival of GBM patients in relation to PXR expression. Kaplan-Meier survival analysis revealed an inverse relationship between PXR expression and patient survival, suggesting that higher levels of PXR may be associated with poorer outcomes in GBM l (**Fig. 4B, left**). Translating this to the *in vitro* model, by analyzing the levels in the U251 GBM line known for being chemo responsive, we observed a downregulation of PXR expression after treatment with TMZ (**Fig.4B, right**). In contrast, through qPCR and immunofluorescence assays, we revealed an increase in PXR levels in all the GBM TMZ-resistant cells after treatment with both TMZ and the combined treatment with *S. Pratensis* (**Fig. 4C-D**). Interestingly, this increase in PXR levels was not associated with an upregulation of its target genes, such as *MDR1, ALDH1A1, and ABCD1*. (**Fig. 4D**).

**Figure 4:**
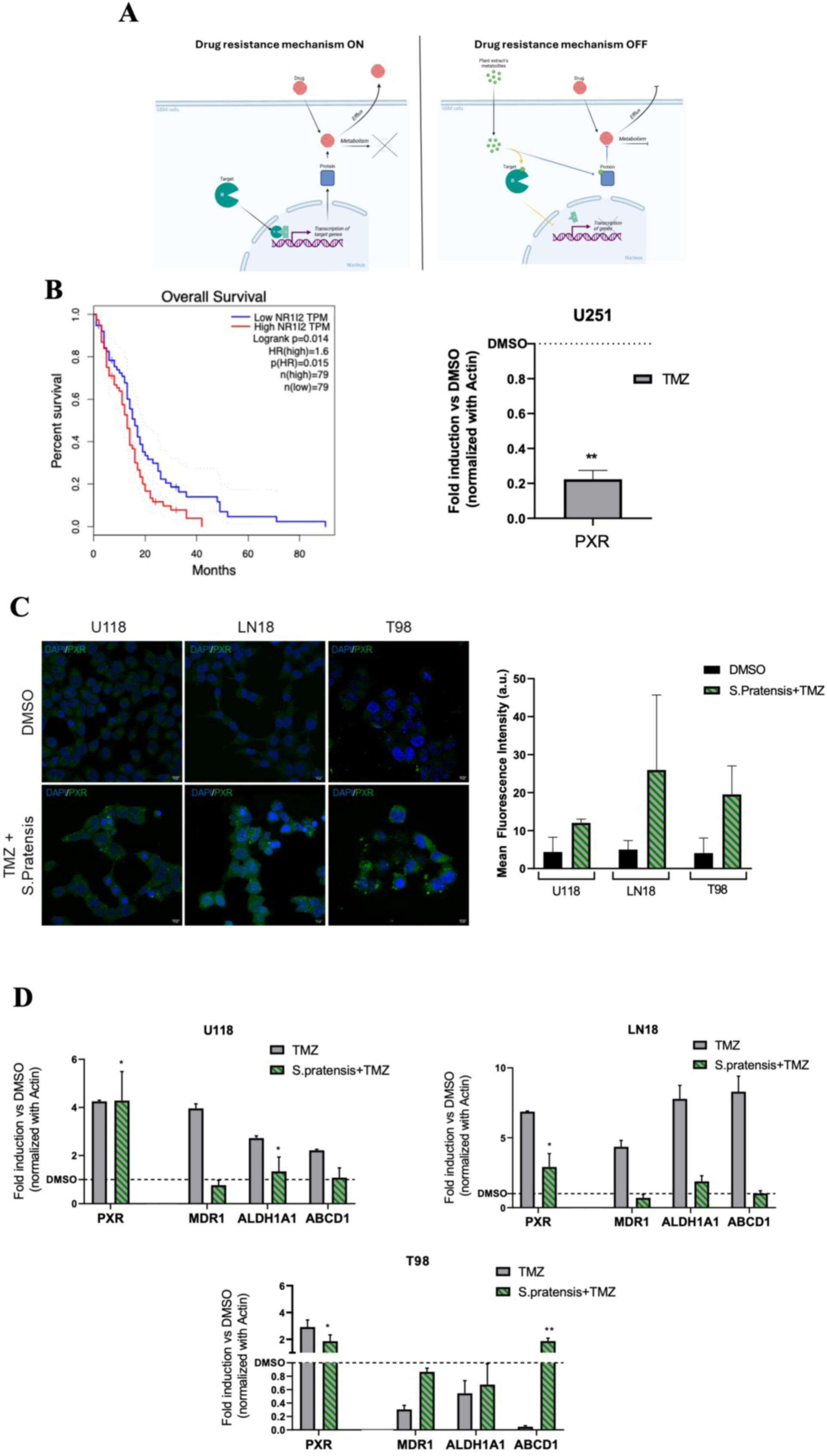
S. Pratensis disrupt GBM drug resistance by modulating PXR activity. **(A)** Graphical representation of the strategy by which *S. Pratensis* modulates drug resistance promoted by PXR. **(B, left side)** Kaplan-Meier survival curves comparing, by RNA-seq clustering, overall survival between patients with high (*red*) and low (*blue*) PXR expression levels. The number of patients in the high-expression and low-expression groups is *n = 79* for each. The hazard ratio (HR) for high PXR expression is 1.6 (*p = 0.015*), suggesting that higher PXR levels are associated with worse overall survival. **(B, right side)** qPCR analysis of PXR expression in U251 cells treated with DMSO (control) or TMZ. Gene expression levels were normalized to β-actin and are presented as fold induction relative to the control. **(C, left side)** Representative immunofluorescence images of cells treated with DMSO (control) or *S. Pratensis* extract combined with TMZ. PXR is stained in green, and nuclei are counterstained with DAPI (blue) **(C, right side).** Fluorescence relative quantification of PXR levels by using ImageJ software. **(D)** qPCR analysis of PXR expression and its target gene levels (*MDR1*, *ALDH1A1,* and *ABCD1)* in cells treated with DMSO (control) or *S. Pratensis* extract combined with TMZ. Gene expression levels were normalized to β-actin and are presented as fold induction relative to the control. (**E**) Representative Western blot analysis showing protein expression levels of PXR (50 kDa), ALDH1A1 (56 kDa), and MDR1 (170 kDa) in cells treated with DMSO (vehicle control), SPA70, *Succisa Pratensis* extract, Akebia, and Apigenin. Total protein staining is shown as a loading control. (**F**) Densitometric quantification of band intensities is as optical density values normalized to total protein levels. Data are presented as mean ± SD from 3 independent experiments. Statistical significance was assessed using one-way ANOVA (*p < 0.05, **p < 0.01, ***p < 0.001).

To further validate these findings at the protein level, we performed Western blot analysis of PXR and its downstream targets in GBM cells treated with S. Pratensis extract and selected secondary metabolites. Consistent with the transcriptional data, PXR protein levels were increased following treatment with S. Pratensis, apigenin, and Akebia-associated compounds, whereas SPA70 treatment led to a reduction in PXR levels. Notably, despite the increased abundance of PXR, the expression of its canonical downstream targets, including MDR1 and ALDH1A1, was significantly reduced across all treatment conditions compared to control cells. Densitometric analysis confirmed a consistent decrease in MDR1 and ALDH1A1 protein levels even in the presence of elevated PXR expression, indicating a functional impairment of PXR signaling. (**Fig. 4E-F**)

To evaluate the effects driven by PXR inhibition, we used a well-known PXR antagonist, SPA70, a competitive antagonist that binds to the nuclear receptor, preventing coactivator recruitment and suppressing the transcription of PXR-induced detoxification genes. [37]

First, we performed cell counting assays on GBM cells. The combination of the PXR antagonist SPA70 with *S. Pratensis* and TMZ elicited a synergistic effect, greatly amplifying cell death compared to treatments with *S. Pratensis* and TMZ alone (**Fig. 5A**).

**Figure 5:**
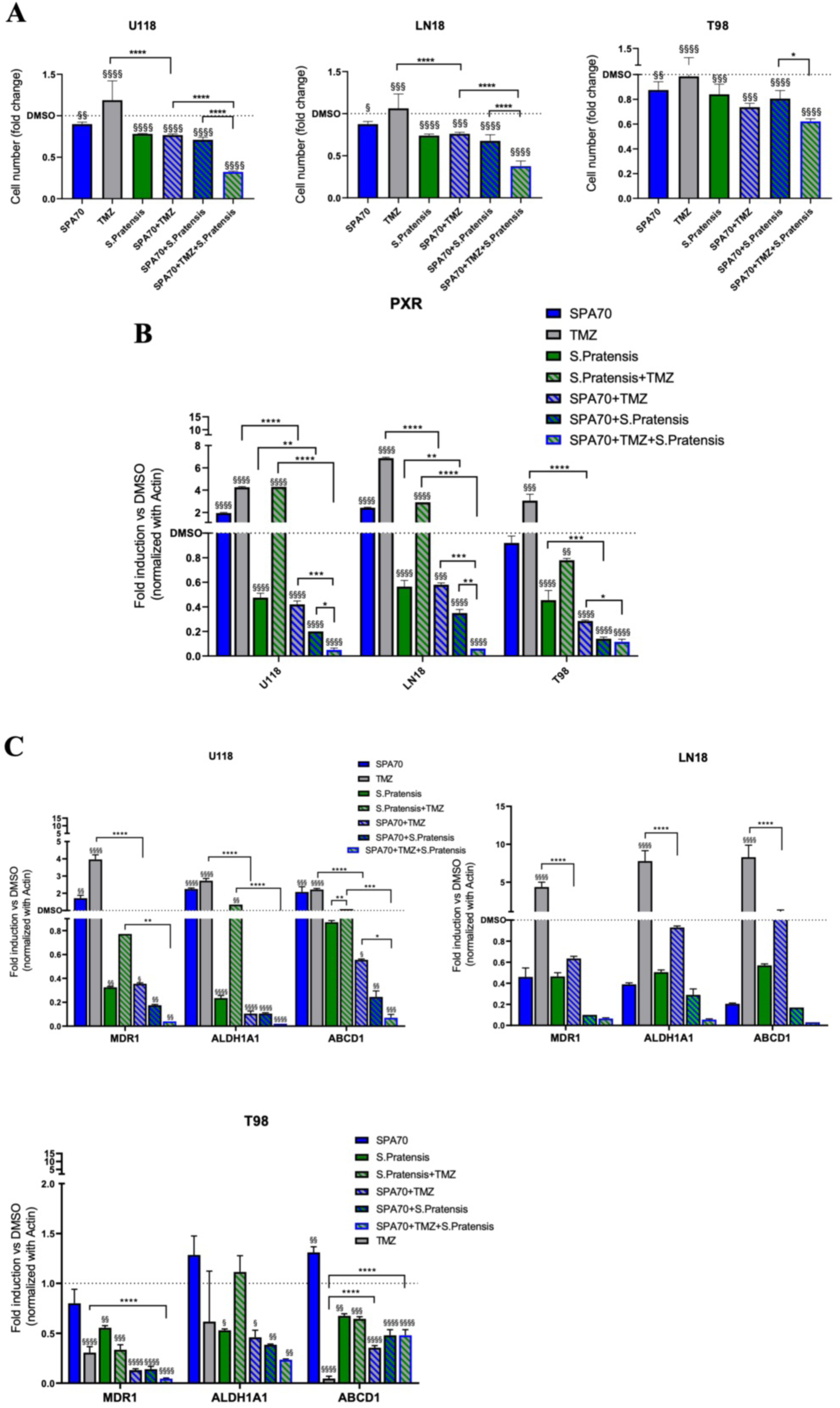
Combined inhibition of PXR by SPA70 and S. pratensis enhances TMZ-induced cytotoxicity and downregulates PXR-mediated efflux pathways in GBM cells. **(A)** Quantification of proliferation assay on the three GBM cells treated with PXR inhibitor (SPA70), TMZ, or *S. Pratensis* whole extract, the double combination (SPA70+TMZ and SPA70+*S. Pratensis*) and the triple combination (SPA70+TMZ+*S. Pratensis*) **(B**) PXR expression level by qPCR analysis in all the GBM cells treated with PXR inhibitor (SPA70), TMZ, or *S. Pratensis* whole extract, the double combination (SPA70+TMZ and SPA70+*S. Pratensis*) and the triple combination (SPA70+TMZ+*S. Pratensis*). Gene expression levels were normalized to β-actin and are presented as fold of induction (FOI) relative to the control. (**C**) qPCR analysis of PXR target gene levels (*MDR1*, *ALDH1A1,* and *ABCD1)* in cells treated with DMSO (control) or with PXR inhibitor (SPA70), TMZ, or *S. Pratensis* whole extract, the double combination (SPA70+TMZ and SPA70+*S. Pratensis*) and the triple combination (SPA70+TMZ+*S. Pratensis*). Gene expression levels were normalized to β-actin and are presented as fold induction relative to the DMSO control. Data represents the mean ± standard deviation (SD) from three independent experiments (*N* = 3). Statistical significance was determined using a t-test (*p < 0.05).

Additionally, we examined the effects of each treatment on PXR and on its target genes. Remarkably, when the PXR antagonist was combined with *S. Pratensis and* TMZ, we observed a reduction not only in PXR expression but also in the expression of its target genes. This suggests a genuine impairment of the efflux system. (**Fig 5B-C**).

To further validate the antitumoral effect of the combinatory treatment, we performed molecular assays to investigate the activation of apoptosis pathways. Specifically, we assessed the expression levels of key components of both the extrinsic TRAIL/DR-5 axis and the intrinsic mitochondrial pathway, focusing on pivotal members of the BCL-2 family, the pro-apoptotic proteins BAX and BAD, and the anti-apoptotic protein BCL-2. (**Fig 6 A-B)**

**Figure 6:**
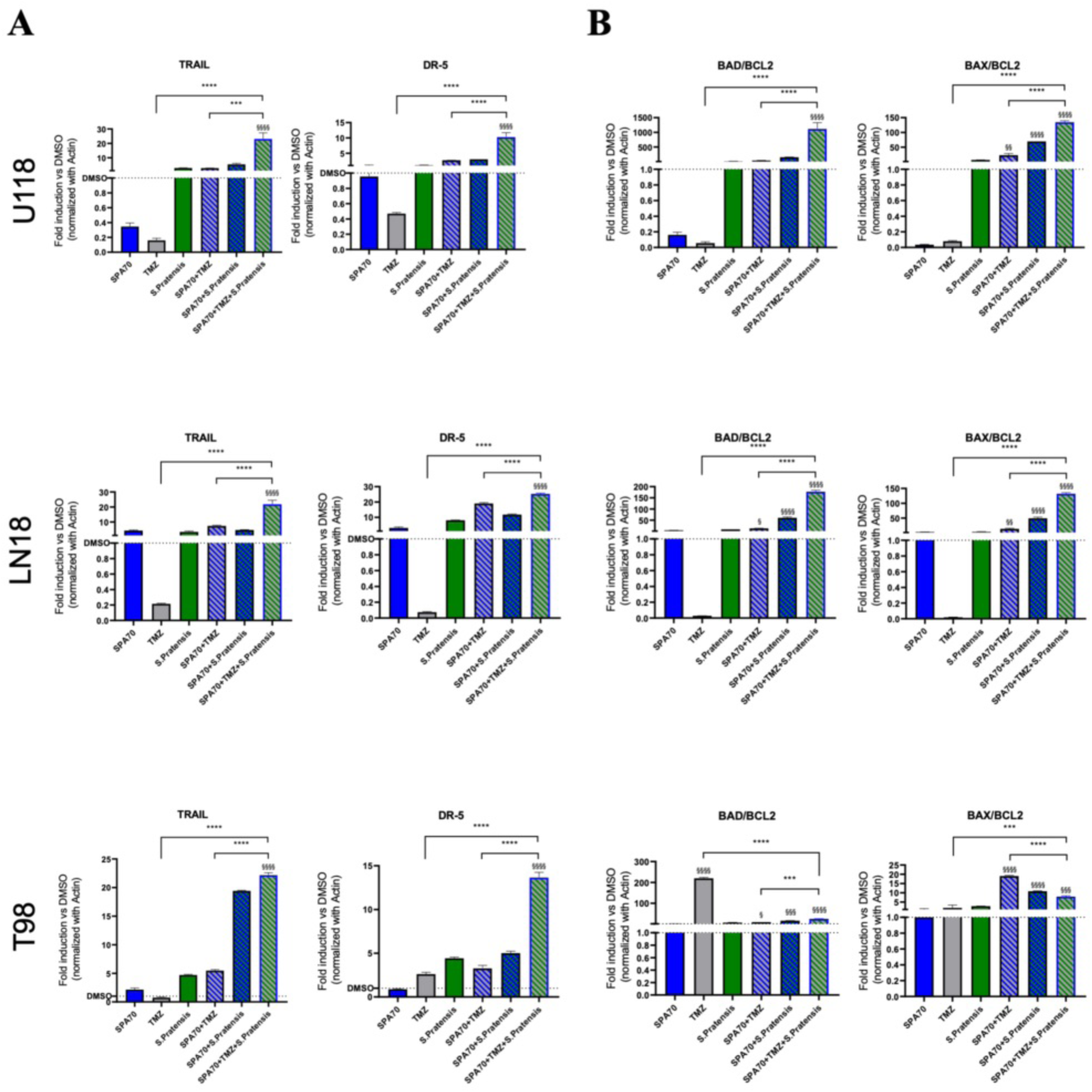
*S. Pratensis* exerts a Pro-Apoptotic effect on Glioma. **(A)** Q- PCR analysis of the extrinsic apoptotic pathway in U118, LN18, and T98 cell lines, expressed by TRAIL and DR5 gene levels. Gene expression data were normalized to β-actin and expressed as FOI to the control condition **(B)**. Evaluation of the intrinsic mitochondrial apoptotic pathway through the expression profiling of the BCL-2 gene family. The activation state of this pathway was represented by the BAD–BAX/BCL2 ratio, reflecting the balance between pro- and anti-apoptotic factors. Gene expression levels were normalized to β-actin and reported as FOI.

We observed a wide death extrinsic TRAIL/DR5 axis activation when cells were treated with the triple combination of SPA70, TMZ, and *S. Pratensis*. The *TRAIL* and *DR-5* expression were highly upregulated in all GBMs compared to the untreated ones or to the single treatment with TMZ (**Fig. 6A**).

Nevertheless, the mitochondrial apoptotic pathway was activated in the U118 and LN18 cells but not in the T98. Specifically, the BCL-2 family balance, indicated by *BAD/BCL2* and *BAX/BCL2* ratio, was highly upregulated by all the single treatments, but notably observed a notable synergistic effect was observed by the triple combination treatment. (**Fig 6B**)

### Apigenin-derived metabolites target PXR and recapitulate chemo sensitization effects

After demonstrating that *S. pratensis* extract combined with TMZ inhibits resistant GBM cell growth through PXR modulation (Fig. 4D, 5A-C), we aimed to identify the bioactive compounds able to bind the receptor ligand-binding domain (LBD) and disrupt downstream gene activation.

UPLC-HRMS analysis of *S. pratensis* leaf extracts identified 21 abundant metabolites, including phenylpropanoids, caffeic acid derivatives, the C-glycosylated flavones apigenin and luteolin, secoiridoid glucosides (oleoside, swertiamarin, gentiopicroside, and sweroside), and the saponin Akebia saponin D (Fig. 7A, Table 2) [14]. To identify potential PXR inhibitors, we performed flexible docking analyses using the LBD coordinates as the docking grid (Fig. 7B, Table 2). For each ligand, affinity values, chemical structures, PubChem CID, and chromatographic retention times were recorded. We focused on the three most abundant metabolites: Akebia saponin D (CID: 14284436), chlorogenic acid (CID: 1794427), and apigenin-C-hexosyl-O-(caffeoyl) hexoside (CID: 44257791). Chlorogenic acid and Akebia saponin D showed affinities of −7.76 and −7.18 kcal/mol, respectively, but displayed distinct binding poses. Chlorogenic acid, owing to its smaller size, docked within the PXR-LBD pocket, whereas the larger Akebia saponin D preferentially localized outside the pocket. Apigenin-C-hexosyl-O-(caffeoyl) hexoside showed the highest affinity (−10.67 kcal/mol) and fitted efficiently within the binding pocket, interacting with amino acids known to stabilize PXR ligands [38].

**Figure 7:**
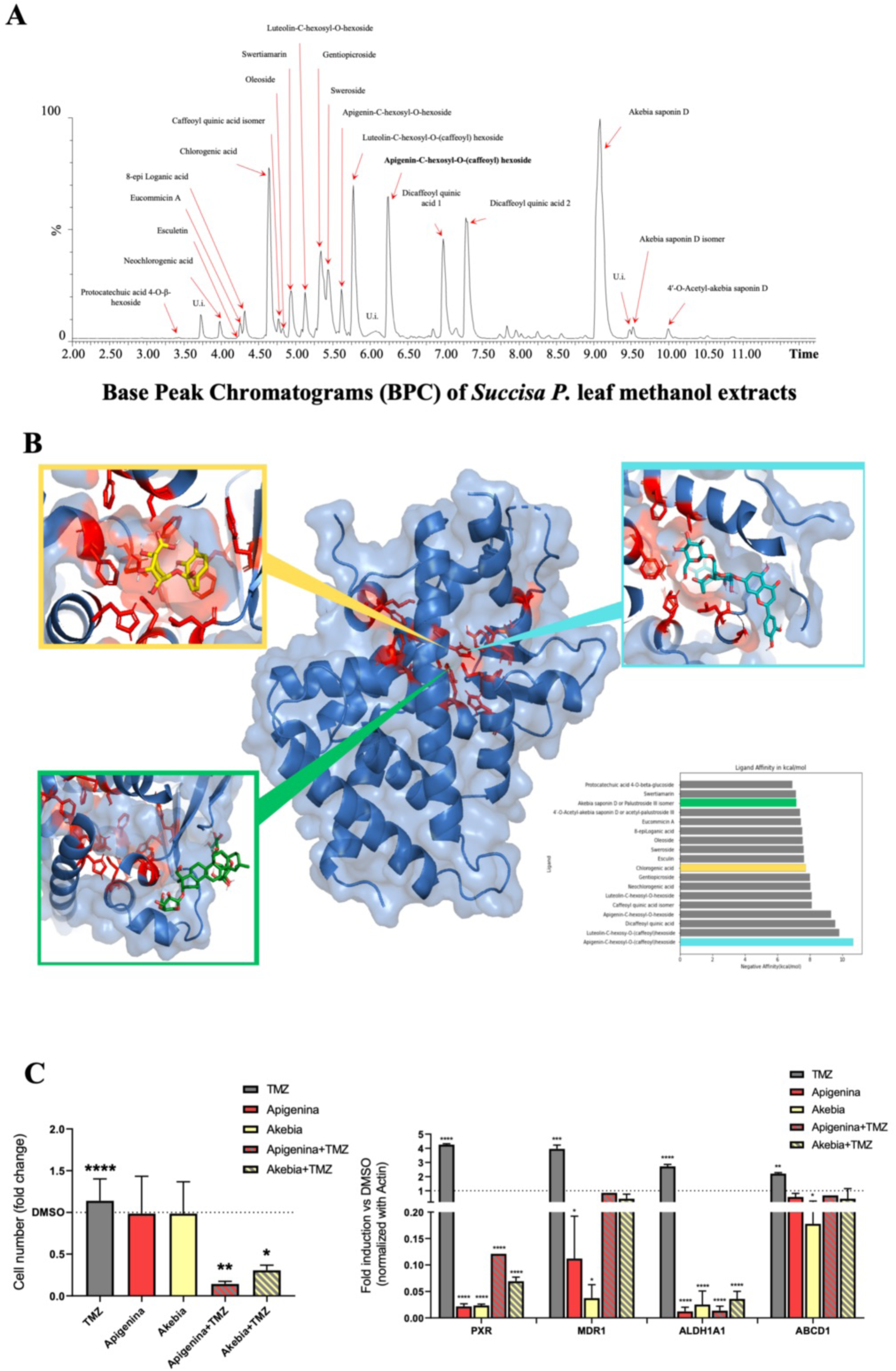
S. Pratensis metabolites replicate the Extract’s Anti-Glioblastoma Action. **(A)** Complete *S. Pratensis* plant extract Chromatogram and related secondary metabolites. **(B)** 3D representation of the PXR receptor and poses of the three most abundant metabolites in the chromatogram detected in *S. Pratensis*. Representation of PXR with highlighted amino acids known to stabilize ligands within the pocket. The poses of Akebia saponin D ligand, Chlorogenic acid, the Apigenin-C-hexosyl-O-(caffeoyl)hexoside, are reported in the green, yellow, and light blue boxes, respectively, with relation to the geometry of the PXR receptor. The associated molecular docking performances (i.e. the binding affinities) of these metabolites are indicated with the same color in the associated barplot. **(C, left side**) Quantification of proliferation assay on GBM, the graph was expressed as cell number fold change to the control. Data are presented as mean ± standard deviation (SD) from three independent experiments (N=3). (C, right side) qPCR of PXR and PXR target gene levels (*MDR1*, *ALDH1A1,* and *ABCD1)* in U118 cells treated with single secondary metabolites (apigenin and akebia), expression levels were normalized to β-actin and are plotted as FOI to the control.

After identifying candidate PXR-LBD inhibitors, we investigated whether individual metabolites could reproduce the effects of the whole *S. pratensis* extract. U118 cells were treated for 24 h with apigenin-C-hexosyl-O-(caffeoyl) hexoside and Akebia saponin D, alone or combined with TMZ. Although the metabolites alone showed no cytotoxicity, they significantly sensitized cells to TMZ, resulting in reduced proliferation (Fig. 7C, left). At the molecular level, treatment with the metabolites, alone or combined with TMZ, significantly downregulated PXR expression together with its target genes MDR1 and ALDH1A1 (Fig. 7C, right). Their antitumoral activity was further supported by apoptosis analyses, which revealed significant activation of the intrinsic mitochondrial pathway (Fig. 8A), whereas no effects were detected on the extrinsic TRAIL/DR5 axis (Fig. 8B-C). These findings suggest that the selected secondary metabolites primarily induce cell death through activation of mitochondrial apoptosis.

**Figure 8:**
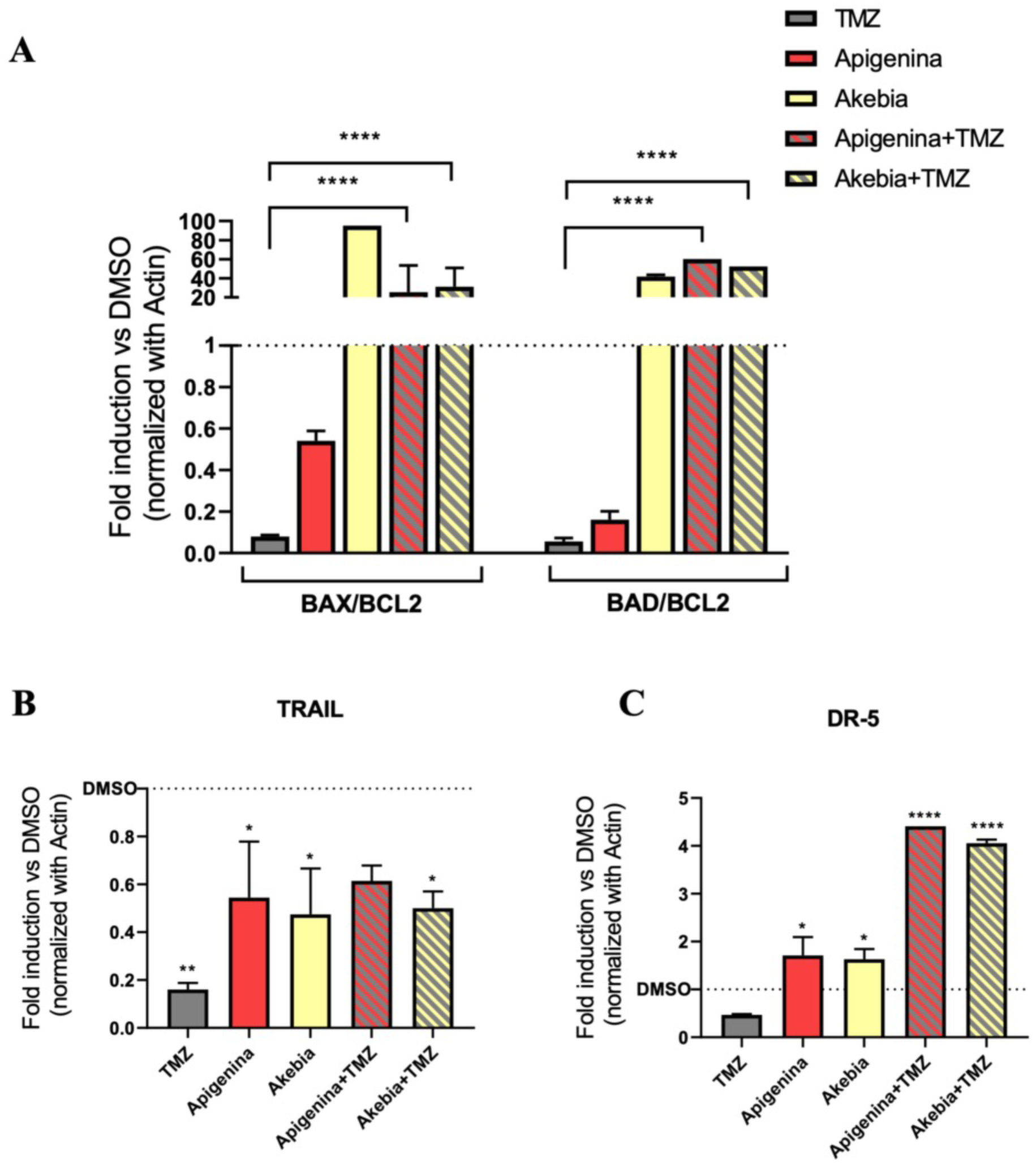
Secondary metabolites from S. pratensis induce GBM cell death predominantly through activation of the intrinsic mitochondrial apoptosis pathway. (**A-C**) Gene expression analysis of intrinsic and extrinsic apoptotic pathways (BCL-2 family and TRAIL axis) was performed by qPCR, normalized to β-actin, and reported as FOI to the control.

## Discussion

GBM is among the most aggressive and deadly adult brain tumors. Despite multimodal treatment with surgery, radiotherapy, and temozolomide (TMZ), it remains highly resistant to therapy and prone to recurrence, highlighting the need for novel strategies able to overcome chemoresistance and improve clinical outcomes. [1–6, 29–30]

The present study investigated natural plant extracts in combination with standard chemotherapy to enhance treatment efficacy. In our previous work, we identified several phytoextracts with anti-GBM activity, with *S. pratensis* showing the strongest effects by inhibiting proliferation, inducing cell-cycle arrest, reducing migration, and promoting MET.

In addition, *S. pratensis* altered GBM metabolism, causing depletion of glycolytic, TCA cycle, and pentose phosphate pathway intermediates, consistent with impaired energy production and anabolic fluxes. Reduced fructose-1,6-bisphosphate may reflect both diminished glycolysis and loss of pyruvate kinase activation, explaining the concomitant decrease in lactate. [33–34] Nucleotide, amino acid, and isoprenoid synthesis were broadly suppressed, whereas GSH and folate one-carbon metabolism increased. Although GSH and GSSG levels rose, the reduced GSH/GSSG ratio indicated a more oxidizing intracellular environment. The decrease in oxidative PPP intermediates together with increased folate metabolites suggests compensatory activation of alternative NADPH-producing pathways, insufficient to restore redox balance. [35–36,38]

Overall, these findings indicate that *S. pratensis* induces oxidative stress and disrupts anabolic metabolism, thereby inhibiting GBM growth and migration. [31–32,38]. Because redox homeostasis and nucleotide metabolism are closely linked to responses to alkylating agents, such metabolic stress may also affect therapy resistance. [37–38]

The upregulation of folate metabolism correlated with increased TMZ sensitization. [44–46] TMZ induces DNA methylation, activating mismatch repair and ultimately causing DNA damage. [10,37] Increased folate intermediates may support nucleotide synthesis and DNA repair attempts; however, in the presence of TMZ-induced lesions, these processes become ineffective, resulting in metabolic exhaustion and apoptosis. [35–36] Moreover, folate metabolism contributes to S-adenosylmethionine production and DNA methylation, potentially influencing MGMT expression and TMZ sensitivity. [35–36,40] This metabolic rewiring likely increases the burden on cancer cells and enhances therapeutic vulnerability.

Proteomic and pathway analyses further highlighted the nuclear receptor PXR as a central regulator of drug resistance. [41–44] Given its established role in xenobiotic metabolism and efflux transporter regulation, PXR represents a critical component of the cellular response to chemotherapy. [16,41–44]

We observed that *S. pratensis*, alone or combined with TMZ, increased PXR expression in resistant GBM cells. However, this was not accompanied by upregulation of canonical target genes such as MDR1, ALDH1A1, and ABCD1. This uncoupling between receptor expression and transcriptional output suggests functional impairment of PXR activity rather than simple downregulation.

A possible explanation lies in the structural properties of secondary metabolites present in *S. pratensis*. Flavonoid derivatives, including apigenin-like compounds, are strong candidates for direct PXR modulation. Their size and planar structure allow interaction with the large and flexible ligand-binding domain (LBD), but likely in a non-productive manner. [16,41,44] Their hydrogen-bonding pattern may fail to stabilize the active receptor conformation and correctly position helix 12 (AF-2), which is required for co-activator recruitment. [16,44] As a result, PXR may remain ligand-bound but transcriptionally inactive, reducing the expression of genes involved in drug resistance.

Although *S. pratensis* is a complex mixture of bioactive compounds, our findings suggest that its chemosensitizing effects are largely attributable to a limited subset of metabolites. Apigenin-derived flavonoids and saponin-associated molecules emerge as key contributors capable of impairing PXR signaling and reducing drug-resistance gene expression. While the whole extract exerts broader antitumoral effects, likely through synergistic interactions, this compositional complexity limits direct clinical translation.

From a translational perspective, whole plant extracts present challenges including compositional variability, poor standardization, and limited pharmacokinetic control. These issues are particularly relevant in GBM, where efficacy depends on BBB penetration. [7,29,30] Large, highly polar molecules, such as many saponins, are unlikely to reach therapeutic concentrations in the brain. In contrast, smaller and structurally defined metabolites such as apigenin derivatives represent more promising candidates for optimization and delivery. These observations support a model in which *S. pratensis* serves primarily as a source of bioactive scaffolds rather than a directly translatable therapeutic agent.

This mechanism is conceptually consistent with the selective PXR antagonist SPA70, which binds the LBD and prevents co-activator recruitment. Whereas SPA70 stabilizes an inactive receptor conformation, apigenin-like flavonoids may induce a less stable but still non-permissive state, resulting in partial or context-dependent antagonism. The enhanced cytotoxicity observed with SPA70 co-treatment further supports the relevance of PXR inhibition.

In contrast, larger metabolites such as the triterpenoid saponin Akebia saponin D are unlikely to act as direct ligands of the canonical PXR binding pocket because of their size and polarity. Docking analyses suggest preferential localization outside the core LBD cavity. Nevertheless, they may contribute indirectly through membrane interactions, signaling pathway modulation, or biotransformation into smaller aglycones capable of interacting with nuclear receptors.

Together, these findings support a model in which *S. pratensis* exerts multi-layered modulation of PXR activity. Small flavonoid-like molecules may directly interfere with receptor activation, while larger saponin-like compounds contribute through indirect or complementary mechanisms. This combinatorial effect may explain the uncoupling between PXR expression and transcriptional activity, leading to reduced activation of efflux systems and enhanced TMZ sensitivity.

Importantly, the whole extract demonstrated stronger antitumoral effects than individual metabolites, likely because of synergistic interactions among different compound classes. While isolated metabolites mainly activated the intrinsic apoptotic pathway, the complete extract also triggered the extrinsic TRAIL-mediated pathway, resulting in a stronger and more consistent induction of cell death.

Overall, *S. pratensis* and its bioactive metabolites exert a profound antitumor effect on GBM cells by disrupting metabolic pathways and attenuating drug-resistance mechanisms. By suppressing glycolysis, the TCA cycle, and nucleotide biosynthesis while enhancing oxidative stress, *S. pratensis* reduces proliferative and migratory capacity. In parallel, modulation of PXR activity appears to play a key role in overcoming chemoresistance.

In summary, our findings indicate that *S. pratensis*-derived metabolites can functionally impair PXR signaling, reducing the expression of drug-resistance genes and increasing TMZ sensitivity. These results identify PXR as a potential therapeutic target and support the use of natural compounds as adjuvant strategies for GBM treatment. Future studies are needed to validate these mechanisms in vivo and to explore the role of PXR modulation in GBM stem-like cells and tumor recurrence.

## Declarations

## Ethical Statement

As this study did not involve human or animal subjects, ethical approval was not required. We declare that no breach of ethical rules was made during the preparation and publication of the study

## Patient consent

Not applicable, as the study did not involve human participants or patient data.

## Consent for publication and conflict of Interest

All authors have read and approved the final version of the manuscript, declare that they have no conflict of interest and specify that no commercial company participated in or contributed to the preparation of the manuscript

## Availability of Data

The research group makes the data available to reviewers or other authors who will request the corresponding author for the data or access to the original analysis files. The identification data for the differentially expressed proteins (DEPs) detected by LC–ESI–MS/MS are hosted in the UNIMI Dataverse repository and downloadable at https://doi.org/10.13130/RD_UNIMI/ME5M6X. All specifications regarding configurations for docking carried out are available at: https://github.com/IBCBC-bioRex/succisa_pratensis_docking.git

## Author Contributions

Conceptualization: **A.G. and A.L.D**.; Methodology: **F.S., F.P., S.M., S.R., M.N., C.P., C.G., and M.M**.; Formal analysis: **F.G. and D.G**.; Investigation: **M.C., B.G., and D.C**.; Data curation: **A.G., A.L.D., D.G., M.B., D.C, B.G., and M.C.;** Writing original draft: **A.G., A.L.D., and F.S.;** Writing review & editing: **G.B., A.G., D. G., and A.L.D**.; Project administration: **A.L.D.;** Funding acquisition: **G.B. D.G. and A.L.D.** All authors have read and agreed to the published version of the manuscript.

## Acknowledgments

The authors are also grateful to Lorena Bonaldi for their advice and administrative support. TMT-LC-MS runs were performed at UNITECH OMICs, an advanced mass spectrometry core facility established by the Università degli Studi di Milano.

## Additional Materials

### Proteomics analysis

Differential proteomics analysis by liquid chromatography–mass spectrometry using TMT (tandem mass tag) isobaric labeling and sample fractionation. Samples were analyzed using a Dionex Ultimate 3000 nano-LC system (Sunnyvale, CA, USA) coupled to an Orbitrap Fusion Tribrid Mass Spectrometer (Thermo Scientific, Bremen, Germany). Data were processed using MaxQuant software (Max Planck Institute, Martinsried, Germany) for protein identification and Perseus software (Max Planck Institute) for statistical analysis of variations, expressed as log_2_ fold change between the two experimental groups. Statistical significance was assessed using ANOVA with false discovery rate correction (p-value < 0.05), followed by Tukey’s post-hoc test for pairwise comparisons.

Pathway analysis was performed using Ingenuity Pathway Analysis software (Qiagen, Hilden, Germany). The “Core Analysis” function was used to interpret the data by analyzing biological processes, canonical pathways, and diseases & biofunctions enriched with differentially regulated proteins. Subsequently, the “Comparison Analysis” function was applied to visualize and identify significantly altered proteins or upstream regulators across experimental conditions. p-values were calculated using a right-tailed Fisher’s exact test. A Fisher’s exact test p-value < 0.05 was considered statistically significant. The activation z-score was used to predict the activation or inhibition of pathways, functions, or regulators.

### Metabolite extraction from cell culture and LC-MS metabolic profiling

The GBM cells were plated in 6-well plates with the above-described culture medium, and after 24h, it was replaced with complete fresh medium in the presence or the absence of S. pratensis at 5mg/ml, then incubated for 24h. Metabolite extraction for LC-MS analysis was performed as described previously [I]. Briefly, cells were rinsed with NaCl 0.9% and then quenched with an ice-cold solution of 70:30 acetonitrile: water. Plates were placed at −80 °C for 10 min and then collected by scraping and sonicated twice for 5 s for five pulses at 70% power. Samples were centrifuged at 12000xg for 10 min and supernatant aqueous phases were collected in a glass insert and dried in a centrifugal vacuum concentrator (Concentrator plus/Vacufuge plus, Eppendorf, Hamburg, Germany) at 30 °C for about 2.5 h. Samples were then resuspended with 150 μL of H2O before analysis. LC separation was performed using an Agilent 1290 Infinity UHPLC system and an Infinity Lab Poroshell 120 PFP column (2.1 × 100 mm, 2.7 μm; Agilent Technologies). Mobile phase A was water with 0.1% formic acid. Mobile phase B was acetonitrile with 0.1% formic acid. The injection volume was 10 μL and LC gradient conditions were: 0 min: 100% A; 2 min: 100% A; 4 min: 99% A; 10 min: 98% A; 11 min: 70% A; 15 min: 70% A; 16 min: 100% A with 2 min of post-run. Flow rate was 0.2 ml/ min, and the column temperature was 35 ◦C. MS detection was performed using an Agilent 6550 iFunnel Q-TOF mass spectrometer with Dual JetStream source operating in negative ionization mode. MS parameters were: gas temp: 285 ◦C; gas flow: 14 l/min; nebulizer pressure: 45 psig; sheath gas temp: 330 ◦C; sheath gas flow: 12 l/min; VCap: 3700 V; Fragmentor: 175 V; Skimmer: 65 V; Octopole RF: 750 V. Active reference mass correction was done through a second nebulizer using masses with m/z: 112.9855 and 1033.9881. Data were acquired from m/z 60–1050. Data analysis and isotopic natural abundance correction were performed with MassHunter ProFinder and MassHunter VistaFlux software (Agilent) as described in [IX].

### Molecular Docking

Molecular docking is a computational technique that predicts how ligands bind to receptor proteins and estimates the strength of these interactions. It models atomic-level interactions between small molecules and target proteins, offering insights into their biochemical mechanisms. As a structure-based method, it relies on high-resolution 3D protein structures obtained through X-ray crystallography, NMR spectroscopy, or Cryo-EM. [II] [III] [IV] For this study, we selected the PDB file 7N2A as a 3D structure related to PXR, from the RCSB Protein Data Bank [V], which is an LBD of human PXR bound to compound 2. To prepare the PXR receptor for docking, we removed compound 2 and water molecules and adjusted the atomic charges. Thereafter, we searched within the PubChem database [VI for structures related to the metabolites detected in the chromatogram of *S. Pratensis.* We then picked the SMILES (Simplified Molecular Input Line Entry System) sequence for each metabolite and converted the sequences into PDB files using the NovoPro Bioscience Inc. tool (https://www.novoprolabs.com/tools/smiles2pdb). Subsequently, we performed molecular docking using AutoDock [VII, a popular molecular docking software developed by the Scripps Research Institute, able to perform both rigid and flexible docking simulations. This software uses a Lamarckian genetic algorithm [VIII] to optimize the positioning of ligands within the binding site of a receptor and incorporates various scoring functions to assess the binding affinity between ligands and receptors. We performed a flexible docking analysis, with a grid set close to the Ligand binding domain (LBD), and evaluated the best poses for each possible ligand, both in terms of affinity and in terms of positioning relative to the PXR. We analyzed the final 3D results using PyMOL (http://www.pymol.org/pymol). All specifications regarding configurations for docking carried out are available at: https://github.com/IBCBC-bioRex/succisa_pratensis_docking.git

### Reaction Activity Scores

To properly assess the possible impact of the proteomics data on metabolism, it is possible to use Reaction Activity Scores (RAS) [IX]. These features are originally derived by mapping the gene expression data to a set of metabolic reactions, organized in a metabolic network model, by means of the Gene Protein Reaction (GPR) rules. These rules describe the relationship between genes and reactions and among genes possibly associated with the same reaction. More in detail, the GPRs represent logical formulas describing how gene products concur to catalyze a given reaction, using Boolean operators. The AND operator is used when distinct genes encode different subunits of the same enzyme, whereas the OR operator is used for genes encoding isoforms of the same enzyme. These AND and OR operators can be combined to describe more complex scenarios involving both isoforms and subunits. The RAS scores are calculated by substituting the mRNA abundances in the corresponding GPRs. Then, the logical expressions are solved by taking the minimum transcript level value when multiple genes are joined by an AND operator, and the sum of their values when multiple genes are joined by an OR operator. In the case of GPRs combining both operators, we respected their standard precedence. Using the same logic, it is possible to map proteomics data to the corresponding set of metabolic reactions by substituting the proteomics abundances in the corresponding GPRs. Using the RAS scores, starting from a set of proteomics profiles, we can generate a set of metabolic profiles, each one consisting of a selected number of RAS scores, each specific to a metabolic reaction.

### Reaction Propensity Scores

To integrate metabolomics data on the metabolic network, we used specific features called Reaction Propensity Scores (RPS). These scores are computed as follows:

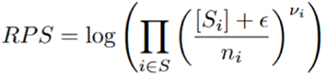

where S_i_ is the concentration of substrate i, nu_i_ is the stoichiometric coefficient of substrate i, n_i_ is the number of reactions in which the substrate i is involved, and eps is a small quantity to avoid log 0. This equation was computed considering that, according to the mass action law, the rate of any chemical reaction is indeed proportional to the product of the concentrations. This assumption holds if the substrate is in significant excess over the enzyme constant. If the reaction is reversible, we define the net RPS as the difference between its forward and backward RPSs.

### The metabolic network model and map

To properly analyze the proteomics and metabolomics differences between the two biological conditions, we used the recently published metabolic network model ENGRO2 [19]. ENGRO2 contains 469 reactions, 395 metabolites, and 497 genes. For this model, 351 model reactions are associated with a gene-protein-reaction (GPR) rule. More in detail, there are 200 single-gene GPRs, 98 OR-expression, 17 AND-expression, and 29 complex rules (i.e., logical expression with both AND and OR operator). With this model, we can study the possible implications of proteomics and metabolomics on a selected number of metabolic pathways mainly involved in human central carbon and essential amino acids metabolism. Moreover, for this model, a manually curated map is available for mapping statistically significant differences between the RAS and RPS of two groups of samples. For this model, we also have a graphical representation designed to enhance clarity and interpretability. More details about the model can be found in the associated publications.

### Mapping RAS and RPS on the metabolic map

We mapped RAS and RPS onto the metabolic network using Marea4Galaxy (26), a Galaxy tool that statistically compares RAS and RPS between two sample groups across all metabolic reactions and visualizes differences on a metabolic map. The tool supports both datasets for integrated multi-omics analysis, displaying RAS differences on reaction arrow bodies and RPS differences on arrowheads. For reversible reactions, only the relevant directional arrowhead is colored based on the net RPS. Further model details are available in the associated publications. [IX]

## References

1. Agnihotri S, Burrell KE, Wolf A, Jalali S, Hawkins C, Rutka JT, et al. Glioblastoma, a Brief Review of History, Molecular Genetics, Animal Models and Novel Therapeutic Strategies. Arch Immunol Ther Exp (Warsz). 2013 Feb 7;61(1):25–41.

2. Ostrom QT, Price M, Neff C, Cioffi G, Waite KA, Kruchko C, et al. CBTRUS Statistical Report: Primary Brain and Other Central Nervous System Tumors Diagnosed in the United States in 2015–2019. Neuro Oncol. 2022 Oct 5;24(Supplement_5):v1–95.

3. Yalamarty SSK, Filipczak N, Li X, Subhan MA, Parveen F, Ataide JA, et al. Mechanisms of Resistance and Current Treatment Options for Glioblastoma Multiforme (GBM). Cancers (Basel). 2023 Apr 1;15(7):2116.

4. Stupp R, Mason WP, van den Bent MJ, Weller M, Fisher B, Taphoorn MJB, et al. Radiotherapy plus Concomitant and Adjuvant Temozolomide for Glioblastoma. New England Journal of Medicine. 2005 Mar 10;352(10):987–96.

5. Maher EA, Furnari FB, Bachoo RM, Rowitch DH, Louis DN, Cavenee WK, et al. Malignant glioma: genetics and biology of a grave matter. Genes Dev. 2001 Jun 1;15(11):1311–33.

6. McAleavey PG, Walls GM, Chalmers AJ. Radiotherapy-drug combinations in the treatment of glioblastoma: a brief review. CNS Oncol. 2022 Jun 30;11(2).

7. de Gooijer MC, Kemper EM, Buil LCM, Çitirikkaya CH, Buckle T, Beijnen JH, et al. ATP-binding cassette transporters restrict drug delivery and efficacy against brain tumors even when blood-brain barrier integrity is lost. Cell Rep Med. 2021 Jan;2(1):100184.

8. Chernov AN, Alaverdian DA, Galimova ES, Renieri A, Frullanti E, Meloni I, et al. The Phenomenon of Multidrug Resistance in Glioblastomas. Hematol Oncol Stem Cell Ther. 2022 Apr;15(2):1–7.

9. Amawi H, Hammad AM, Hall FS, Hussein N, Rataan AO, Mrayyan A, et al. Revisiting strategies to target ABC transporter-mediated drug resistance in CNS cancer. Cancer Biol Med. 2025 Sep 29;1–23.

10. Zhang J, F.G. Stevens M, D. Bradshaw T. Temozolomide: Mechanisms of Action, Repair and Resistance. Curr Mol Pharmacol. 2012 Jan 1;5(1):102–14.

11. Ju HQ, Lin JF, Tian T, Xie D, Xu RH. NADPH homeostasis in cancer: functions, mechanisms and therapeutic implications. Signal Transduct Target Ther. 2020 Oct 7;5(1):231.

12. Cena H, Labra M. Biodiversity and planetary health: a call for integrated action. The Lancet. 2024 May;403(10440):1985–6.

13. Cragg GM, Newman DJ. Plants as a source of anti-cancer agents. J Ethnopharmacol. 2005 Aug 22;100(1–2):72–9.

14. Giammona A, Commisso M, Bonanomi M, Remedia S, Avesani L, Porro D, et al. A Novel Strategy for Glioblastoma Treatment by Natural Bioactive Molecules Showed a Highly Effective Anti-Cancer Potential. Nutrients. 2024 Jul 23;16(15):2389.

15. Witkowska-Banaszczak E, Krajka-Kuźniak V, Papierska K. The effect of luteolin 7-glucoside, apigenin 7-glucoside and Succisa pratensis extracts on NF-κB activation and α-amylase activity in HepG2 cells. Acta Biochim Pol. 2020 Mar 4;

16. Lehmann JM, McKee DD, Watson MA, Willson TM, Moore JT, Kliewer SA. The human orphan nuclear receptor PXR is activated by compounds that regulate CYP3A4 gene expression and cause drug interactions. Journal of Clinical Investigation. 1998 Sep 1;102(5):1016–23.

17. Morrison C, Weterings E, Mahadevan D, Sanan A, Weinand M, Stea B. Expression Levels of RAD51 Inversely Correlate with Survival of Glioblastoma Patients. Cancers (Basel). 2021 Oct 26;13(21):5358.

18. Planque C, Rajabi F, Grillet F, Finetti P, Bertucci F, Gironella M, et al. Pregnane X-receptor promotes stem cell-mediated colon cancer relapse. Oncotarget. 2016 Aug 30;7(35):56558–73.

19. Mareboina M, Bakhl K, Agioti S, Yee NS, Georgakopoulos-Soares I, Zaravinos A. Comprehensive Analysis of Granzymes and Perforin Family Genes in Multiple Cancers. Biomedicines. 2025 Feb 7;13(2):408.

20. Weber GL, Parat MO, Binder ZA, Gallia GL, Riggins GJ. Abrogation of PIK3CA or PIK3R1 reduces proliferation, migration, and invasion in glioblastoma multiforme cells. Oncotarget. 2011 Nov 30;2(11):833–49.

21. Sciorra VA, Sanchez MA, Kunibe A, Wurmser AE. Suppression of Glioma Progression by Egln3. PLoS One. 2012 Aug 8;7(8):e40053.

22. Chou V, Pearse R V., Aylward AJ, Ashour N, Taga M, Terzioglu G, et al. INPP5D regulates inflammasome activation in human microglia. Nat Commun. 2023 Nov 29;14(1):7552.

23. Gu QY, Liu YX, Wang JL, Huang XL, Li RN, Linghu H. LLGL2 Inhibits Ovarian Cancer Metastasis by Regulating Cytoskeleton Remodeling via ACTN1. Cancers (Basel). 2023 Dec 18;15(24):5880.

24. Huang J, Zhang T, Li H, Li Z, Yin S, Liu Y, et al. Ubiquitination-Dependent LLGL2 Degradation Drives Colorectal Cancer Progression via *THBS3* mRNA Stabilization. Advanced Science. 2025 Oct 6;12(39).

25. Sun P, Yan F, Fang W, Zhao J, Chen H, Ma X, et al. MDM4 contributes to the increased risk of glioma susceptibility in Han Chinese population. Sci Rep. 2018 Jul 23;8(1):11093.

26. Zhang C, Yang M, Li Y, Tang S, Sun X. FOXA1 is upregulated in glioma and promotes proliferation as well as cell cycle through regulation of cyclin D1 expression. Cancer Manag Res. 2018 Sep;Volume 10:3283–93.

27. Lin W, Wang YM, Chai SC, Lv L, Zheng J, Wu J, et al. SPA70 is a potent antagonist of human pregnane X receptor. Nat Commun. 2017 Sep 29;8(1):741.

28. Bansard L, Laconde G, Delfosse V, Huet T, Ayeul M, Rigal E, et al. Targeting pregnane X receptor with a potent agonist-based PROTAC to delay colon cancer relapse. Oncogenesis. 2025 Aug 30;14(1):34.

29. Wu W, Klockow JL, Zhang M, Lafortune F, Chang E, Jin L, et al. Glioblastoma multiforme (GBM): An overview of current therapies and mechanisms of resistance. Pharmacol Res. 2021 Sep;171:105780.

30. Singh S, Dey D, Barik D, Mohapatra I, Kim S, Sharma M, et al. Glioblastoma at the crossroads: current understanding and future therapeutic horizons. Signal Transduct Target Ther. 2025 Jul 9;10(1):213.

31. D’Aprile S, Denaro S, Gervasi A, Vicario N, Parenti R. Targeting metabolic reprogramming in glioblastoma as a new strategy to overcome therapy resistance. Front Cell Dev Biol. 2025 Feb 26;13.

32. Bernhard C, Reita D, Martin S, Entz-Werle N, Dontenwill M. Glioblastoma Metabolism: Insights and Therapeutic Strategies. Int J Mol Sci. 2023 May 23;24(11):9137.

33. Wang M, Huang X, Zhang D, Liu Y, Liu P. The role of fructose-1,6-bisphosphatase 1 on regulating the cancer progression and drug resistance. Discover Oncology. 2025 Mar 18;16(1):346.

34. Rho H, Hay N. Protein lactylation in cancer: mechanisms and potential therapeutic implications. Exp Mol Med. 2025 Mar 24;57(3):545–53.

35. Newman AC, Maddocks ODK. One-carbon metabolism in cancer. Br J Cancer. 2017 Jun 4;116(12):1499–504.

36. Zarou MM, Vazquez A, Vignir Helgason G. Folate metabolism: a re-emerging therapeutic target in haematological cancers. Leukemia. 2021 Jun 11;35(6):1539–51.

37. Li H, Wu Y, Chen Y, Lv J, Qu C, Mei T, et al. Overcoming temozolomide resistance in glioma: recent advances and mechanistic insights. Acta Neuropathol Commun. 2025 Jun 5;13(1):126.

38. Ciccarese F, Ciminale V. Escaping Death: Mitochondrial Redox Homeostasis in Cancer Cells. Front Oncol. 2017 Jun 9;7.

39. Muller IB, Lin M, Jonge R, Will N, López-Navarro B, Laken C van der, et al. Methotrexate Provokes Disparate Folate Metabolism Gene Expression and Alternative Splicing in Ex Vivo Monocytes and GM-CSF- and M-CSF-Polarized Macrophages. Int J Mol Sci. 2023 Jun 1;24(11):9641.

40. Hervouet E, Debien E, Campion L, Charbord J, Menanteau J, Vallette FM, et al. Folate Supplementation Limits the Aggressiveness of Glioma via the Remethylation of DNA Repeats Element and Genes Governing Apoptosis and Proliferation. Clinical Cancer Research. 2009 May 15;15(10):3519–29.

41. Xing Y, Yan J, Niu Y. PXR: a center of transcriptional regulation in cancer. Acta Pharm Sin B. 2020 Feb;10(2):197–206.

42. Pondugula SR, Mani S. Pregnane xenobiotic receptor in cancer pathogenesis and therapeutic response. Cancer Lett. 2013 Jan;328(1):1–9.

43. Niu X, Wu T, Li G, Gu X, Tian Y, Cui H. Insights into the critical role of the PXR in preventing carcinogenesis and chemotherapeutic drug resistance. Int J Biol Sci. 2022;18(2):742–59.

44. Willson TM, Kliewer SA. Pxr, car and drug metabolism. Nat Rev Drug Discov. 2002 Apr;1(4):259–66.

## References

I. Sahoo RN, Pattanaik S, Pattnaik G, Mallick S, Mohapatra R. Review on the use of Molecular Docking as the First Line Tool in Drug Discovery and Development. Indian J Pharm Sci. 2022;84(5).

II. Meng XY, Zhang HX, Mezei M, Cui M. Molecular Docking: A Powerful Approach for Structure-Based Drug Discovery. Current Computer Aided-Drug Design. 2011 Jun 1;7(2):146–57.

III. Agu PC, Afiukwa CA, Orji OU, Ezeh EM, Ofoke IH, Ogbu CO, et al. Molecular docking as a tool for the discovery of molecular targets of nutraceuticals in diseases management. Sci Rep. 2023 Aug 17;13(1):13398.

IV. Berman HM. The Protein Data Bank. Nucleic Acids Res. 2000 Jan 1;28(1):235–42.

V. Kim S, Chen J, Cheng T, Gindulyte A, He J, He S, et al. PubChem 2025 update. Nucleic Acids Res. 2025 Jan 6;53(D1):D1516–25.

VI. Morris GM, Huey R, Lindstrom W, Sanner MF, Belew RK, Goodsell DS, et al. AutoDock4 and AutoDockTools4: Automated docking with selective receptor flexibility. J Comput Chem. 2009 Dec 27;30(16):2785–91.

VII. Fuhrmann J, Rurainski A, Lenhof H, Neumann D. A new Lamarckian genetic algorithm for flexible ligand-receptor docking. J Comput Chem. 2010 Jul 15;31(9):1911–8

VIII. Damiani C, Rovida L, Maspero D, Sala I, Rosato L, Di Filippo M, et al. MaREA4Galaxy: Metabolic reaction enrichment analysis and visualization of RNA-seq data within Galaxy. Comput Struct Biotechnol J. 2020;18:993–9.

IX. Di Filippo M, Pescini D, Galuzzi BG, Bonanomi M, Gaglio D, Mangano E, et al. INTEGRATE: Model-based multi-omics data integration to characterize multi-level metabolic regulation. PLoS Comput Biol. 2022 Feb 7;18(2):e1009337.

