## Supplementary figures and images for "Disrupting Pregnane X Receptor Signaling Overcomes Temozolomide Resistance in Glioblastoma via Succisa pratensis–Derived Metabolites"

### supplementary info 1

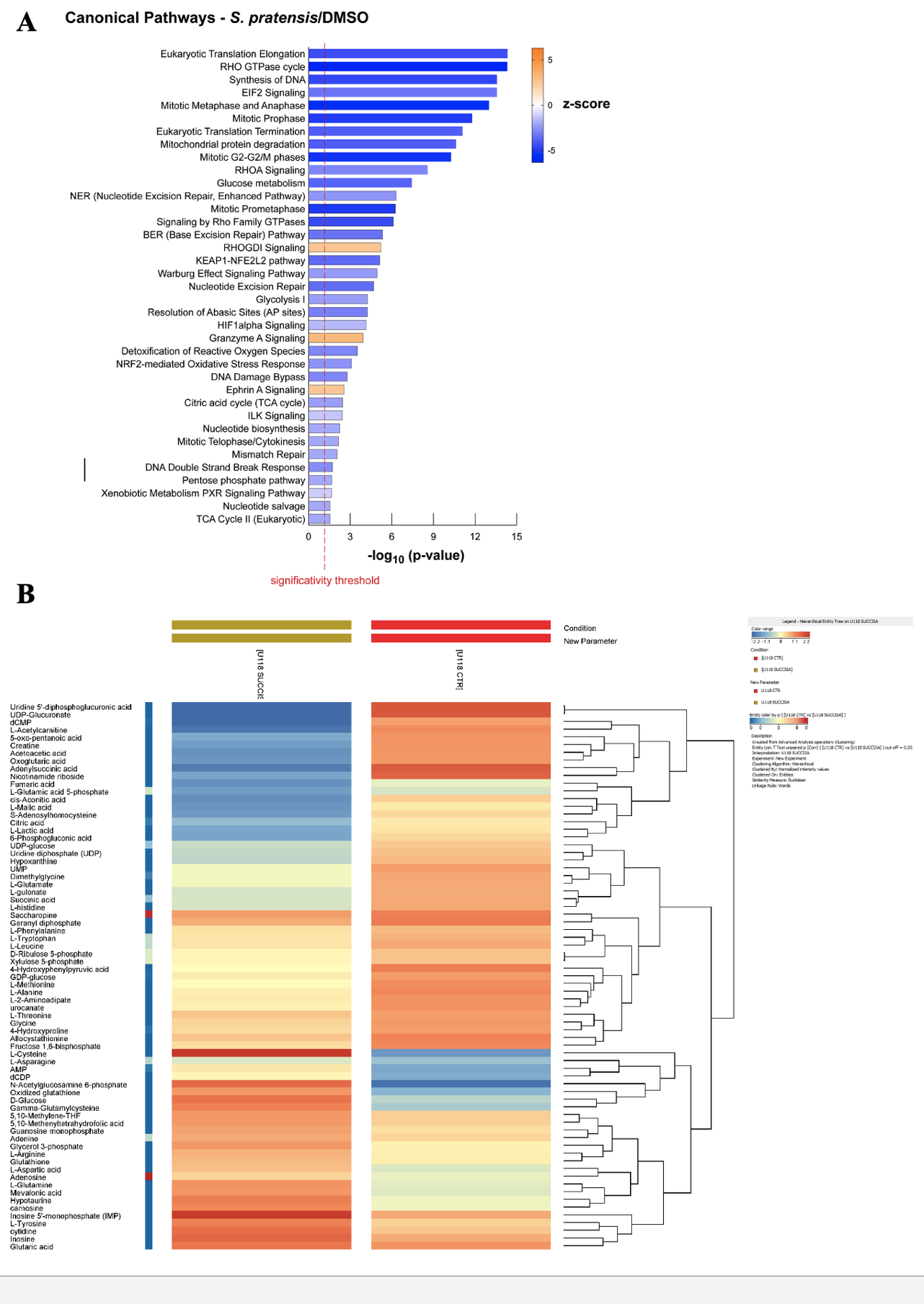

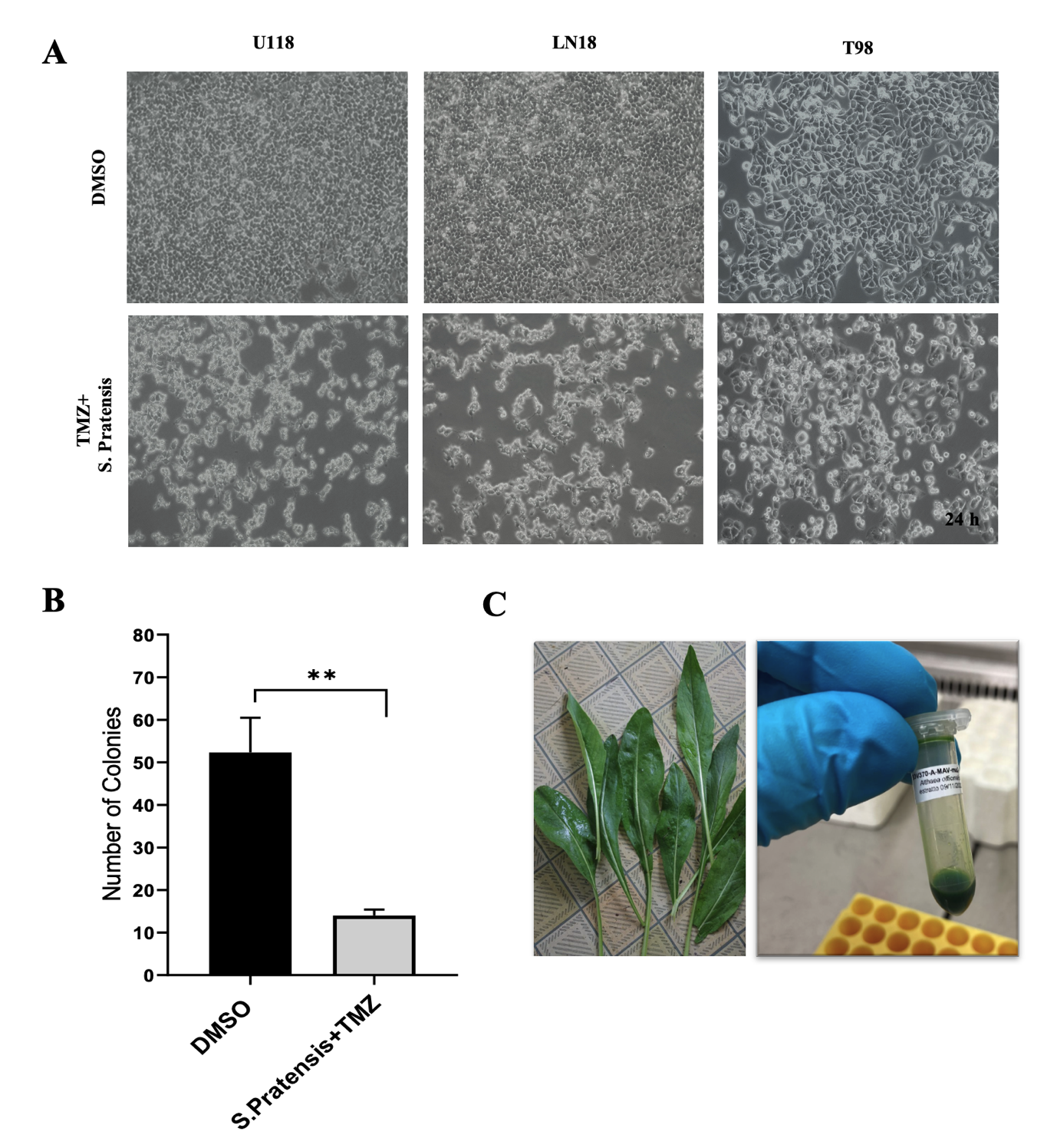
